# Exponential increase in QTL detection with increased sample size

**DOI:** 10.1101/2023.01.27.525982

**Authors:** Apurva S. Chitre, Oksana Polesskaya, Daniel Munro, Riyan Cheng, Pejman Mohammadi, Katie Holl, Jianjun Gao, Hannah Bimschleger, Angel Garcia Martinez, Anthony George, Alexander F. Gileta, Aidan Horvath, Alesa Hughson, Keita Ishiwari, Christopher P. King, Alexander Lamparelli, Cassandra L. Versaggi, Connor Martin, Celine L. St. Pierre, Jordan A. Tripi, Jerry B. Richards, Tengfei Wang, Hao Chen, Shelly B. Flagel, Paul Meyer, Terry E. Robinson, Leah C. Solberg Woods, Abraham A. Palmer

## Abstract

Power analyses are often used to determine the number of animals required for a genome wide association analysis (GWAS). These analyses are typically intended to estimate the sample size needed for at least one locus to exceed a genome-wide significance threshold. A related question that is less commonly considered is the number of significant loci that will be discovered with a given sample size. We used simulations based on a real dataset that consisted of 3,173 male and female adult N/NIH heterogeneous stock (HS) rats to explore the relationship between sample size and the number of significant loci discovered. Our simulations examined the number of loci identified in sub-samples of the full dataset. The sub-sampling analysis was conducted for four traits with low (0.15 ± 0.03), medium (0.31 ± 0.03 and 0.36 ± 0.03) and high (0.46 ± 0.03) SNP-based heritabilities. For each trait, we sub-sampled the data 100 times at different sample sizes (500, 1,000, 1,500, 2,000, and 2,500). We observed an exponential increase in the number of significant loci with larger sample sizes. Our results are consistent with similar observations in human GWAS and imply that future rodent GWAS should use sample sizes that are significantly larger than those needed to obtain a single significant result.

## Introduction

Genome wide association studies (GWAS) in both humans and rodents have been extremely successful in understanding the genetics of quantitative traits. Outbred rodent populations such as Heterogeneous stock (HS) rats, Diversity Outbred (DO) mice, and Advanced Intercross Lines (AIL) have proven to be an invaluable resource for genetic mapping studies. The success of these outbred rodent strains can be attributed to the ability to provide high resolution QTL mapping (Solberg Woods and Palmer 2019). With each generation of recombination, the number of markers and independent tests increases, which in turn increases the threshold for statistical significance. In comparison to an F_2_ cross, outbred rodent populations offer better resolution for mapping QTLs (Solberg Woods 2014; Gonzales and Palmer 2014). Inbred rodent strains such as the Hybrid Rat Diversity panels (HRDP), Hybrid Mouse Diversity Panels (HMDP) and Recombinant Inbred (RI) strains (such as the BXD and CC panels) have also been successfully employed for mapping studies (Williams and Williams 2017). However, the sample size involving these panels is usually limited by the number of strains available in the population. QTL mapping studies are not limited to rodent populations. These genetic studies are also conducted in zebrafish (Kwon et al. 2019), fruit flies (Wangler et al. 2017) and plants such as *Arabidopsis thaliana* (Togninalli et al. 2020).

In GWAS studies power is defined as the likelihood of detecting a single significant QTL of a certain effect size. Power analyses are often performed for GWAS studies so that an appropriate sample size can be selected. In general, larger sample sizes increase the power to detect significant loci in humans (Spencer et al. 2009), rodents (Li et al. 2006; Keele et al. 2019), livestock (Wittenburg et al. 2020) and crops (Wang and Xu 2019). Software to perform power analyses has also typically focused on power to detect a single locus given its effect size (Sen et al. 2007; Delongchamp et al. 2018).

In this study, we sought to examine a related question, namely the relationship between sample size and the number of significant loci discovered. We used simulations based on a real dataset that consisted of 3,173 male and female adult N/NIH heterogeneous stock (HS) rats. This dataset is part of our recent publication on the GWAS of obesity related traits in HS rats, which is among the largest rodent GWAS ever performed (Chitre et al. 2020). The dataset in Chitre et al. was collected as part of a large multi-site project focused on genetic analyses of behavioral phenotypes related to drug abuse in HS rats (www.ratgenes.org). We repeatedly subsampled this dataset to determine the number of significant loci that could be identified with various sample sizes.

## Results

The number of significant loci discovered increased exponentially as sample size increased. **Figure 1** shows the average number of significant loci detected for each trait at each sample size. When we ran the analysis with the maximum number of individuals, we detected 28 loci for body weight (h^2^ = 0.46 ± 0.03), 16 loci for body length_Tail (h^2^ = 0.36 ± 0.03), 5 loci for BMI_Tail (h^2^ = 0.31 ± 0.03) and 3 for fasting glucose (h^2^ = 0.15 ± 0.03). As expected, fewer QTLs were discovered with smaller sample sizes. We note the largest increase in the number of QTL detected for body weight, the trait with the highest heritability, with more than a ten-fold increase in detected QTL when the sample size is increased from 500 to 2500. Similar trends are seen for both BMI and fasting glucose.

**Figure 1.**
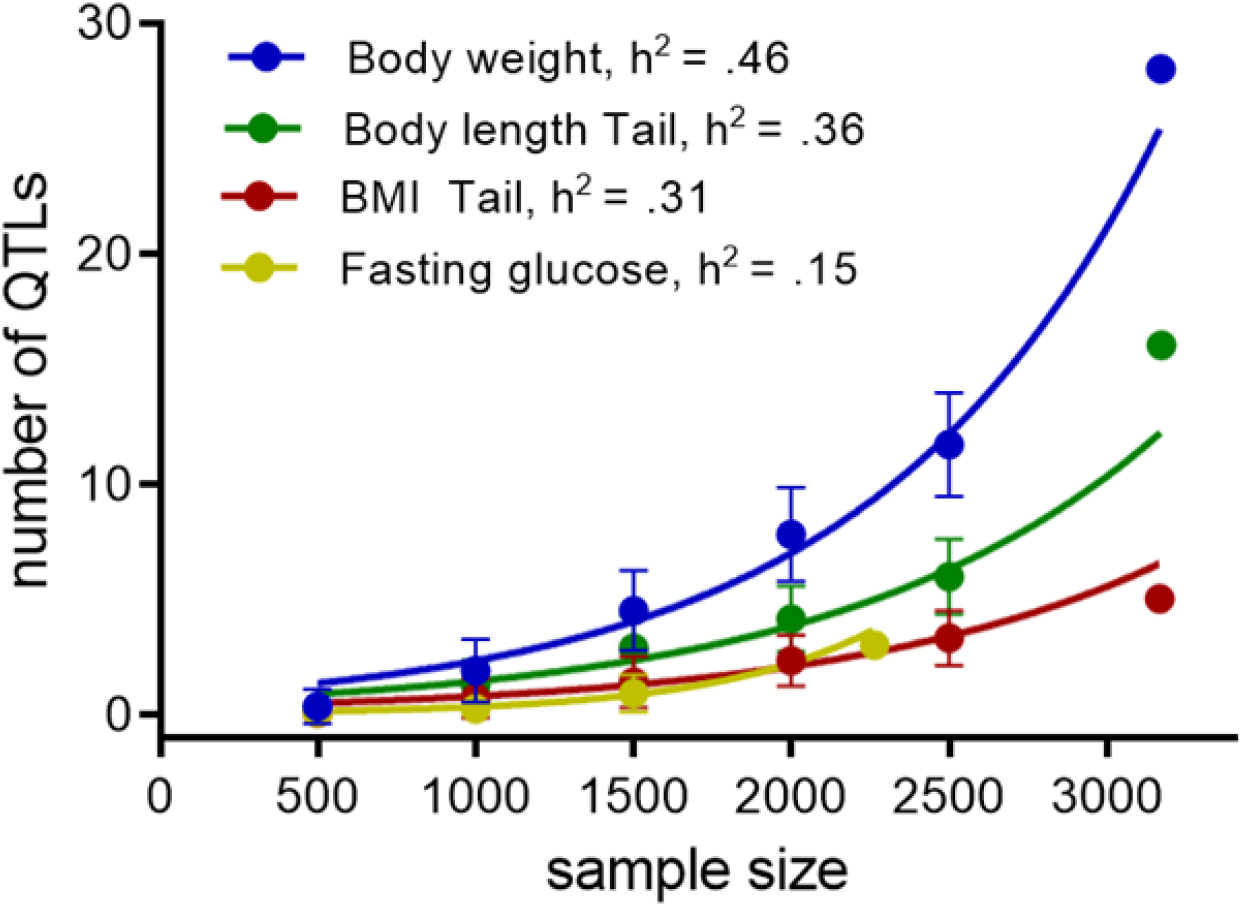
Number of detected QTLs increases with the increase of sample size. Each dot is an average number of QTLs obtained in 100 GWAS, each performed on a randomly selected subset of the actual dataset. Error bars indicate standard deviation. The final point (at ∼3100 animals for body weight, body length_Tail, BMI_Tail and at 2,246 for fasting glucose) used the full dataset and therefore does not include error bars. This simulation was performed on four traits with different heritability: body weight (h^2^ = 0.46 ± 0.03), body length_Tail (h^2^ = 0.36 ± 0.03), BMI_Tail (h^2^ = 0.31 ± 0.03) and fasting glucose (h^2^ = 0.15 ± 0.03).

To determine whether the increase in the number of significant loci was more consistent with a linear or an exponential (log-linear) function, we fitted both models on the data to identify least squares parameters. The two models we defined as

Linear: y = b_0_ x + b_1_ + e,

and

Exponential: y = exp(b_0_ x + b_1_) + e

where, b_0_ and b_1_ are the model parameters, x is the sample size, y is the average number of QTLs and e is the error term.

Since both models have the same number of parameters we compared them in terms of residual sum of squares (RSS) and used bootstrapping to ascertain statistical boundaries of the estimates. We found that an exponential curve fits better; the estimated 95% confidence interval is (0.344, 7.67) for the exponential fit, and (9.57, 217.522) for the linear fit.

We also conducted linkage analysis using haplotypes to confirm that our findings were in agreement with those obtained from GWAS analysis that utilized SNPs. We found that, similar to the GWAS analysis, an exponential increase in the number of QTL identified with increasing sample size for linkage analysis using haplotypes (**Supplementary Figure 1**). We performed this analysis for BMI with tail (SNP *h*^*2*^ = 0.31 ± 0.03) using R/qtl2 (Broman et al. 2019) using a permutation derived threshold of 18.2 LOD at alpha = 0.05. We used the residual sum of squares to compare the linear and exponential models. The RSS values for exponential fit (0.069) are smaller than for linear fit (1.217), suggesting that an exponential curve fits better than a linear curve.

## Discussion

In this study, we used a real dataset to explore the effect of sample size on the number of significant loci identified. This represents a conceptually different approach compared to conventional power analyses, which focus on estimating power to detect at least one genome-wide significant locus. We found an exponential increase in the number of QTL identified with increasing sample size, particularly for body weight, a trait with relatively high heritability. Our results suggest (but do not prove) that our findings would generalize to other similar laboratory populations (HS/Npt, HS-CC, DO, etc.). The results from the haplotype-based linkage mapping analysis also support an exponential increase in the number of QTL identified with increasing sample size for the trait BMI with tail (SNP *h*^*2*^ = 0.31 ± 0.03).

Similar observations in human genetics (Visscher et al. 2012; Sullivan et al. 2018) suggest an initial exponential growth in the number of loci, which is what we have observed, followed by a linear phase when increasing sample size produces a linear increase in the number of significant loci. In the current study, we did not find strong evidence of this linear phase. This could reflect the fact that our sample size, which is still small by the standards of human genetics, was not large enough to get beyond the initial exponential phase. As is the case in human GWAS, the effect size of loci that require larger sample sizes will tend to be smaller than those identified with larger sample sizes, assuming a constant allele frequency. There are several reasons that this dataset was able to identify multiple significant loci despite having a sample size that is smaller than those typically used for human GWAS. First, the effect sizes of alleles discovered in model systems are often much larger than alleles found in humans. The reasons for this are unknown but might include relaxed selection in captive breeding populations, which allows alleles that would have been selected against in a natural population to rise to high frequency. A second reason that smaller sample sizes are sufficient in model systems is that the linkage disequilibrium among SNPs is greater, meaning that fewer tests are performed, thus reducing the multiple testing burden and correspondingly the threshold for significance. The greater LD might also mean that multiple smaller alleles are inherited in blocks that have greater effect sizes. A third advantage of model systems is that they are often created by crossing a small number of inbred strains, meaning that allele frequencies are higher; greater power is always available when alleles are more common. Despite these differences, our observation of exponential growth in the number of significant loci with increasing sample size is very similar to observations in human genetics.

Our result indicates that many previous studies, which have performed power analyses designed to assure that they find a single significant locus, are likely underpowered to find multiple loci that have diminishing effect sizes. Our recommendation is that future studies of complex traits in outbred rodents should use significantly larger sample sizes since they are likely to provide a larger number of findings; this recommendation assumes that the cost of increasing sample size is linear, however in some cases there might be efficiencies of scale that would make the addition of each additional subject less expensive. Fewer studies with larger sample sizes, rather than larger numbers of studies with modest sample sizes might be preferable. Alternatively, multiple traits from separate studies that are genetically correlated might be jointly analyzed, since this can provide some of the advantages of larger sample sizes assuming that certain loci are important for more than one of the traits under study.

## Materials and Methods

The data used in this study are thoroughly described in our recent publication (Chitre et al. 2020). Briefly, phenotypic data on body weight and length (which permit calculation of BMI), fat pad weight, and fasting glucose levels of adiposity traits were collected at three different research sites at multiple ages. We used data from all three sites. Prior to combining data from the three sites, we regressed out the effects of covariates and then performed quantile normalizations within each site and within each sex after which data from all sites and sexes were combined and jointly analyzed to explore the relationship between sample size and the number of significant loci identified.

HS rats used in this study were obtained from the NMcwi:HS colony which was initiated by the NIH in 1984 by interbreeding eight inbred founder strains and were subsequently maintained as an outbred population, making them ideal for fine mapping of genetic loci (Hansen and Spuhler 1984; Solberg Woods and Palmer 2019). Rats were genotyped at 3.4 million autosomal SNPs, however, because there was extensive LD among these SNPs and to reduce computational burden, we used LD pruning (r^2^<0.95) which yielded a set of 128,477 SNPs that were used for all analyses described in this paper.

To determine the number of QTLs detected by different samples sizes, we subsampled data from four phenotypes chosen to have low (0.15 ± 0.03; fasting glucose), medium (0.36 ± 0.030; body length_Tail and .31 ± 0.03; BMI) and high (0.46 ± 0.03; body weight) chip heritabilities (calculated using GCTA). For each dataset, we performed 100 random subsamples in which we retained 500, 1,000, 1,500, 2,000, or 2,500 individuals (for fasting glucose we could not include 2,000 and 2,500 because the total sample size was smaller than 2,000). Thus, we produced 1,300 total subsamples for the three phenotypes. We then performed a GWAS for each subsampled dataset using an automated pipeline based on the LMM software package GEMMA (Zhou and Stephens 2012); we implemented the leave one chromosome out (LOCO) method (Cheng et al. 2013). We have previously shown that an LMM in conjunction with the LOCO methods effectively controls type I error rate (Gonzales et al. 2018; Gileta et al. 2022), meaning that our observations in this study are unlikely to be due to type I errors that can be caused by population structure.

Our pipeline used an algorithm to automatically record the number of significant QTLs in each subsampled dataset. First, we scanned each chromosome to determine if there was at least one SNP that exceeded the threshold of –log_10_(p) > 5.6, which is the threshold used in Chitre et al. 2020. To avoid situations where only a single, presumably anomalous, SNP showed a significant association, we required that at least one other SNP within 0.5 Mb have a p-value that was within 2 –log_10_(p) of the index SNP. If we found a second supporting SNP, we recorded the identification of a QTL for that dataset. Some chromosomes were expected to contain more than one independent QTL, but we were also concerned that we might count a single locus twice. To avoid counting the same locus twice, we excluded all SNPs with r^2^ > 0.4 relative to the just identified index SNP. We then rescanned the chromosome to see if any additional SNPs on this chromosome exceeded the threshold of –log_10_(p) > 5.6. If they did and they were supported by a second SNP within 0.5 Mb that had a p-value that was within 2 – log_10_(p) of the index SNP, we recorded an additional QTL for that dataset. We then repeated these steps as often as needed until no further significant QTLs could be identified on a given chromosome. We then continued this process for all subsequent chromosomes. After scanning the last chromosome, we tabulated the number of QTLs detected for that dataset. We repeated this procedure for each of the 1,300 subsampled datasets. In this way, we determined the number of significant QTLs in 100 possible sub-samplings of each of four traits when using 500, 1,000, 1,500, 2,000, and 2,500 individuals, and in the maximal number of individuals (∼3100 for all traits except fasting glucose).

We performed linkage mapping with haplotypes using R/qtl2 (Broman et al. 2019). We estimated founder haplotypes using the calc_genoprob_fst function with the cohort and founder strain genotypes. We used the scan1perm function to perform 1,000 permutations for establishing the significance threshold. The kinship matrices were derived using the “leave one chromosome out” method with the calc_kinship function. For each sub-sampled dataset for the trait BMI with tail, we performed a genome scan using a linear mixed model with the scan1 function. We used the function find_peaks to identify LOD peaks that exceeded the permutation derived threshold of 18.2.

## Data availability

The data presented in the study are deposited in the UC San Diego Library Digital Collections repository at https://library.ucsd.edu/dc/object/bb9156620z (DOI https://doi.org/10.6075/J0Q240F0).

## Acknowledgements

This work was supported by the National Institute on Drug Abuse (P50 DA037844) and the National Institute of Diabetes and Digestive and Kidney Diseases (R01 DK106386)

## References

Broman, K. W., Gatti, D. M., Simecek, P., Furlotte, N. A., Prins, P., Sen, Ś., Yandell, B. S., & Churchill, G. A. (2019). R/qtl2: Software for Mapping Quantitative Trait Loci with High-Dimensional Data and Multiparent Populations. Genetics, 211(2), 495–502. https://doi.org/10.1534/genetics.118.301595

Cheng, R., Parker, C. C., Abney, M., & Palmer, A. A. (2013). Practical Considerations Regarding the Use of Genotype and Pedigree Data to Model Relatedness in the Context of Genome-Wide Association Studies. G3 Genes Genomes Genetics, 3(10), 1861–1867. https://doi.org/10.1534/g3.113.007948

Chitre AS, Polesskaya O, Holl K, Gao J, Cheng R, Bimschleger H, Garcia Martinez A, George T, Gileta AF, Han W, et al. 2020. Genome□Wide Association Study in 3,173 Outbred Rats Identifies Multiple Loci for Body Weight, Adiposity, and Fasting Glucose. Obesity. 28(10):1964–1973.

Delongchamp R, Faramawi MF, Feingold E, Chung D, Abouelenein S. 2018. The Association between SNPs and a Quantitative Trait: Power Calculation. European Journal of Environment and Public Health [Internet]. [accessed 2020 Dec 28] 2(2). http://www.ejeph.com/article/the-association-between-snps-and-a-quantitative-trait-power-calculation-3925

Gileta, A. F., Fitzpatrick, C. J., Chitre, A. S., St. Pierre, C. L., Joyce, E. V., Maguire, R. J., McLeod, A. M., Gonzales, N. M., Williams, A. E., Morrow, J. D., Robinson, T. E., Flagel, S. B., & Palmer, A. A. (2022). Genetic characterization of outbred Sprague Dawley rats and utility for genome-wide association studies. PLOS Genetics, 18(5), e1010234. https://doi.org/10.1371/journal.pgen.1010234

Gonzales NM, Palmer AA. 2014. Fine-mapping QTLs in advanced intercross lines and other outbred populations. Mamm Genome. 25(7–8):271–292.

Gonzales, N. M., Seo, J., Hernandez Cordero, A. I., St. Pierre, C. L., Gregory, J. S., Distler, M. G., Abney, M., Canzar, S., Lionikas, A., & Palmer, A. A. (2018). Genome wide association analysis in a mouse advanced intercross line. Nature Communications, 9(1), 5162. https://doi.org/10.1038/s41467-018-07642-8

Hansen C, Spuhler K. 1984. Development of the National Institutes of Health genetically heterogeneous rat stock. Alcohol Clin Exp Res. 8(5):477–479.

Keele GR, Crouse WL, Kelada SNP, Valdar W. 2019. Determinants of QTL Mapping Power in the Realized Collaborative Cross. G3. 9(5):1707–1727.

Kwon RY, Watson CJ, Karasik D. 2019. Using zebrafish to study skeletal genomics. Bone. 126:37–50.

Li X, Quigg RJ, Zhou J, Xu S, Masinde G, Mohan S, Baylink DJ. 2006. A critical evaluation of the effect of population size and phenotypic measurement on QTL detection and localization using a large F2 murine mapping population. Genet Mol Biol. 29(1):166–173.

Sen S, Satagopan JM, Broman KW, Churchill GA. 2007. R/qtlDesign: inbred line cross experimental design. Mamm Genome. 18(2):87–93.

Solberg Woods LC. 2014. QTL mapping in outbred populations: successes and challenges. Physiol Genomics. 46(3):81–90.

Solberg Woods LC, Palmer AA. 2019. Using Heterogeneous Stocks for Fine-Mapping Genetically Complex Traits. Methods Mol Biol. 2018:233–247.

Spencer CCA, Su Z, Donnelly P, Marchini J. 2009. Designing Genome-Wide Association Studies: Sample Size, Power, Imputation, and the Choice of Genotyping Chip.Storey JD, editor. PLoS Genet. 5(5):e1000477.

Sullivan PF, Agrawal A, Bulik CM, Andreassen OA, Børglum AD, Breen G, Cichon S, Edenberg HJ, Faraone SV, Gelernter J, et al. 2018. Psychiatric Genomics: An Update and an Agenda. Am J Psychiatry. 175(1):15–27.

Togninalli M, Seren Ü, Freudenthal JA, Monroe JG, Meng D, Nordborg M, Weigel D, Borgwardt K, Korte A, Grimm DG. 2020. AraPheno and the AraGWAS Catalog 2020: a major database update including RNA-Seq and knockout mutation data for Arabidopsis thaliana. Nucleic Acids Res. 48(D1):D1063–D1068.

Visscher PM, Brown MA, McCarthy MI, Yang J. 2012. Five years of GWAS discovery. Am J Hum Genet. 90(1):7–24.

Wang M, Xu S. 2019. Statistical power in genome-wide association studies and quantitative trait locus mapping. Heredity. 123(3):287–306.

Wangler MF, Hu Y, Shulman JM. 2017. Drosophila and genome-wide association studies: a review and resource for the functional dissection of human complex traits. Dis Model Mech. 10(2):77–88.

Williams RW, Williams EG. 2017. Resources for Systems Genetics. In: Schughart K, Williams RW, editors. Systems Genetics [Internet]. Vol. 1488. New York, NY: Springer New York; [accessed 2020 Dec 16]; p. 3–29. http://link.springer.com/10.1007/978-1-4939-6427-7_1

Zhou X, Stephens M. 2012. Genome-wide efficient mixed-model analysis for association studies. Nat Genet. 44(7):821–824.

